# N-glycosylation profiles of the SARS-CoV-2 spike D614G mutant and its ancestral protein characterized by advanced mass spectrometry

**DOI:** 10.1101/2021.07.26.453787

**Authors:** Dongxia Wang, Bin Zhou, Theodore Keppel, Maria Solano, Jakub Baudys, Jason Goldstein, M.G. Finn, Xiaoyu Fan, Asheley P. Chapman, Jonathan L. Bundy, Adrian R. Woolfitt, Sarah Osman, James L. Pirkle, David E. Wentworth, John R. Barr

**Author notes:** To whom correspondence should be addressed. Dongxia Wang, Ph.D., John R. Barr, Ph.D. Person of Contact: Dongxia Wang, Ph.D. **Disclaimer:** The findings and conclusions in this report are those of the author(s) and do not necessarily represent the official position of the Centers for Disease Control and Prevention.

## Abstract

N-glycosylation plays an important role in the structure and function of membrane and secreted proteins. The spike protein on the surface of the severe acute respiratory syndrome coronavirus 2 (SARS-CoV-2), the virus that causes COVID-19, is heavily glycosylated and the major target for developing vaccines, therapeutic drugs and diagnostic tests. The first major SARS-CoV-2 variant carries a D614G substitution in the spike (S-D614G) that has been associated with altered conformation, enhanced ACE2 binding, and increased infectivity and transmission. In this report, we used mass spectrometry techniques to characterize and compare the N-glycosylation of the wild type (S-614D) or variant (S-614G) SARS-CoV-2 spike glycoproteins prepared under identical conditions. The data showed that half of the N-glycosylation sequons changed their distribution of glycans in the S-614G variant. The S-614G variant showed a decrease in the relative abundance of complex-type glycans (up to 45%) and an increase in oligomannose glycans (up to 33%) on all altered sequons. These changes led to a reduction in the overall complexity of the total N-glycosylation profile. All the glycosylation sites with altered patterns were in the spike head while the glycosylation of three sites in the stalk remained unchanged between S-614G and S-614D proteins.

## Introduction

The novel coronavirus SARS-CoV-2 emerged in China in late 2019 and led to the COVID-19^1^ pandemic. Of the major structural proteins encoded by the SARS-CoV-2 genome, the Spike protein (S) has attracted considerable research interest because of the central role it plays in entry into host cells, tissue tropism, species specificity and is the antigenic target of all the approved vaccine candidates to date. Like the S proteins of other related coronaviruses, it is present on the virion as a homotrimer of approximately 180 kDa per protomer, as recently determined by cryogenic electron microscopy (cryo-EM) analysis^2^. Each of these protomers is composed of two subunits. The S1 subunit comprises the head of the molecule that contains the receptor binding domain (RBD) which serves to bind to human cell surface receptor-angiotensin converting enzyme 2 (ACE2)^3^. The S2 domain comprises the distal part of the molecule and mediates fusion of the virus envelope with the host cell membrane. Similar to related viral entry proteins, S has been shown to be extensively glycosylated with 22 N-glycosylation sites present per protomer^2^. The heavy glycosylation can form a glycan shield that functions as a means for the virus to escape antigenic recognition^4, 5^ as well as stabilizing the RBD in a conformation favorable for binding with ACE2^6, 7^. The presence of the glycan shield may also be needed for viral entry, as shown in a recent report where glycosylation was inhibited by chemical and genetic methods^8^.

As the spread of the virus has progressed globally, many sequence variants in the S protein have been observed in clinical isolates using whole genome sequencing. Of these, the substitution of aspartic acid (D) to glycine (G) at residue 614 of the spike protein (S-D614G) has been of highest prevalence, first being detected in early January 2020 in China and Europe and quickly spreading worldwide, being present in over 75% of sequenced samples from infected individuals by late 2020^9, 10^. As of July 2021, nearly all SARS viruses circulating contain S-614G mutation including all of the SARS-CoV-2 variants classified as Variants of Concern (B.1.1.7/Alpha, P.1/Gamma, B1.351/Beta, and B.1.617.2/Delta) and Variants of Interest (B.1.427/Epsilon, B.1.429/Epsilon,B.1.525/Eta, B.1.526/Iota, B.1.617.1/Kappa, B.1.617.3, and P2/Zeta)^11, 12^. Yurkovetskiy and co-workers performed *in vitro* studies in human lung cells comparing S-614D and S-614G variants and found increased infectivity by the S-614G variant^13^. Similar enhanced rates of infection were also observed in a panel of S mutants in studies by Li et al^14^ and in another report using virus like particles (VLPs) expressing S-D614G^15^. Other studies reinforcing these observations found increased rates of viral replication in lung epithelial cells and tissues infected with the S-D614G virus^16, 17^. They also observed increased levels of virus in the upper respiratory tract in hamsters compared to lung tissues. These observations were also independently seen in another report by Zhou and colleagues^18^, in ferrets, hamsters, and a novel mouse ACE2 knock-in model, which showed the S-614G variant had a competitive advantage in replication and transmission.

Cryogenic electron microscopy (Cryo-EM) data reported by Yurkovetskiy et al. indicated that the S-D614G variant significantly favored more “open” conformations compared to the wild type by disrupting interchain contacts stabilized by hydrogen bonding between D614 and residue T859 on an adjacent molecule^13^. These conformations result in the RBD being in an “up” position, facilitating interaction with the ACE2 receptor. The preference for the open conformation in S-D614G was also postulated by Mansbach et al. from molecular dynamic (MD) studies on the variant sequence^19^ and was supported by a recent cryo-EM study of S-D614G^20^. Additionally, molecular dynamics studies have suggested that glycans on N165 and N234 stabilize the open form.^7^ Presumably, this open conformation favors increased RBD interactions with ACE2, and thus greater infectivity.

In recent years, mass spectrometry has emerged as the method of choice for the site specific analysis of protein glycosylation both in global proteomics analyses^21^ and targeted studies on individual or smaller groups of proteins as encountered in virology and vaccinology research^22, 23^. Based on measured molecular size and fragmentation pattern of a glycopeptide, the specific site of glycosylation, glycan composition, and relative abundance of the modified peptide can be determined by mass spectrometry, although more comprehensive analysis is required on the determination of the structure of a modified glycan such as branching, sugar moiety, and linkage^24^. Not surprisingly, the use of this technology for the analysis of the glycan complement of SARS-CoV-2 S protein has resulted in considerable research activity^25–27^. The McLellan group published an initial report describing the glycans of the stabilized trimer construct previously developed in their laboratory, showing heterologous occupancy of all 22 predicted N-glycosites^28^. Subsequently, several other groups have published glycoproteomics analyses performed on individual S1 and S2 constructs^29^ and full-length recombinant S proteins^6, 30^. We employed a glycoproteomics approach based on signature ions triggered electron-transfer dissociation mass spectrometry^31^ to characterize SARS-CoV-1 and −2 spike proteins, and Middle East Respiratory Syndrome (MERS) S protein from multiple expression systems^32^. A common theme observed in all these investigations is that the glycan structures of the S protein is dependent upon both the expression system used to produce the protein, and the structure of the construct (subunit, ectodomain or stabilized ectodomain). These observations have important implications in the use of these reagents for the study of SARS-CoV-2 S protein structure and virus biology.

Given the current predominance of the S-D614G variant and the apparent influence of this modification upon the structure and function of the S protein, it is important to understand how this substitution might influence the glycan complement. Here we use our previously described approach^32^ for analyzing S proteins expressed in a human cell line (HEK 293) for a comparative glycoproteomics study of the S-D614G variant relative to the wild type S. Interestingly, we find that the variant protein, while having a similar glycan profile to the wild type construct at several sites, has significant variations in glycan speciation and quantity at sites both proximal and distal to the substitution.

## Results and Discussion

### Preparation and analysis of recombinant S-614G and S-614D proteins

The glycosylation profiles of recombinant ectodomain of the SARS-CoV-2 S-614G and its progenitor S-614D were examined. To provide accurate comparison, two recombinant proteins were expressed under identical experimental conditions. Constructs of the two isogenic proteins included two widely used substitutions: a double proline mutations at residues 986 and 987 to stabilize the prefusion conformation and amino acids RRAR (position 682-685) were mutated to GSAS to disrupt the furin cleavage site between S1 and S2 subunits ^2^, which aids purification of the whole S ectodomain. Figure 1 is the 1D gel of purified S-614D and S-614G along with the results of an affinity pulldown of the two spike variants with ACE2 and with monoclonal antibodies. The 1D SDS-PAGE gel shows the high purity of both S-614D and S-614G and their binding ability to ACE2 as well as the two monoclonal antibodies^33^ derived from the immunization against the SARS-CoV-2 spike receptor binding domain.

**Figure 1.**
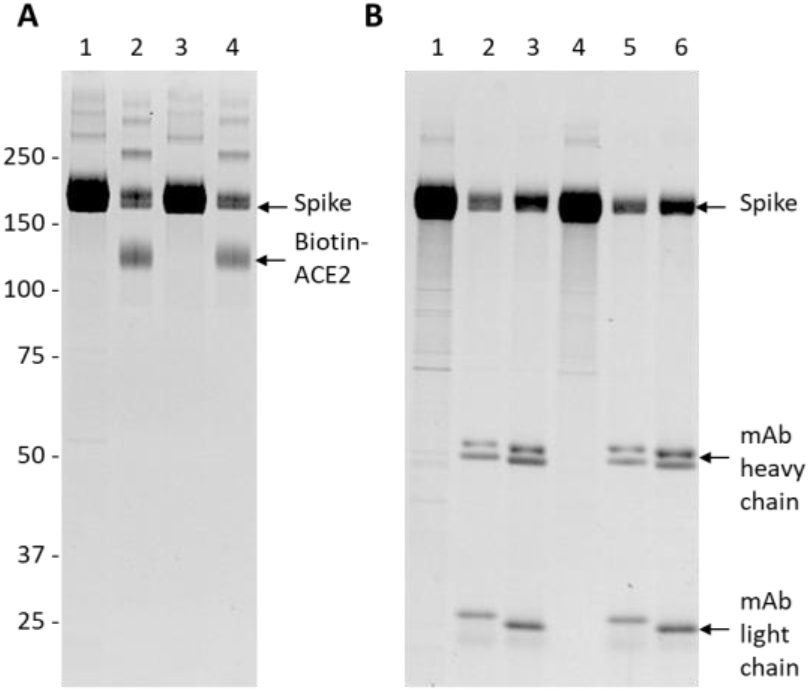
SDS gel analysis of the SARS-CoV-2 S proteins interacting with other proteins. (A) binding of the S to the biotinylated ACE2. lane 1: S-614D; lane 2: S-614D + ACE2; lane 3: S-614G; lane 4: S-614G +ACE2. (B) binding of the S to monoclonal antibodies. Lane 1 – 6: S-614D, S-614D + 3G7, S-614D + 3A2, S-614G, S-614G + 3G7, and S-614G + 3A2. The gels were run under reduced condition and visulized with SYPRO Ruby stains.

The site-specific distribution and abundance of heterogeneous N-linked glycans on 21 of 22 potential sequons were characterized by mass spectrometry analysis of the glycopeptides cleaved by the α-lytic protease or the combination of trypsin and chymotrypsin using the same EThcD instrument parameters described previously (N149, glycosylated peptides were not of sufficient quality for quantification). For direct comparison, the ion intensity of the precursor MS1 peak of each glycopeptide was used to represent the abundance of each individual glycosylation form on a specific sequon of the proteins (Figure 2). The relative abundance of three major types of the N-glycans: high-mannose (HexNAc_2_Hex_>4_X, green), hybrid (3 HexNAc, purple), and complex (> 3 HexNAc, gray), on the N of each sequon of S-614D and S-614G proteins were depicted in the inserted pie charts.

**Figure 2.**
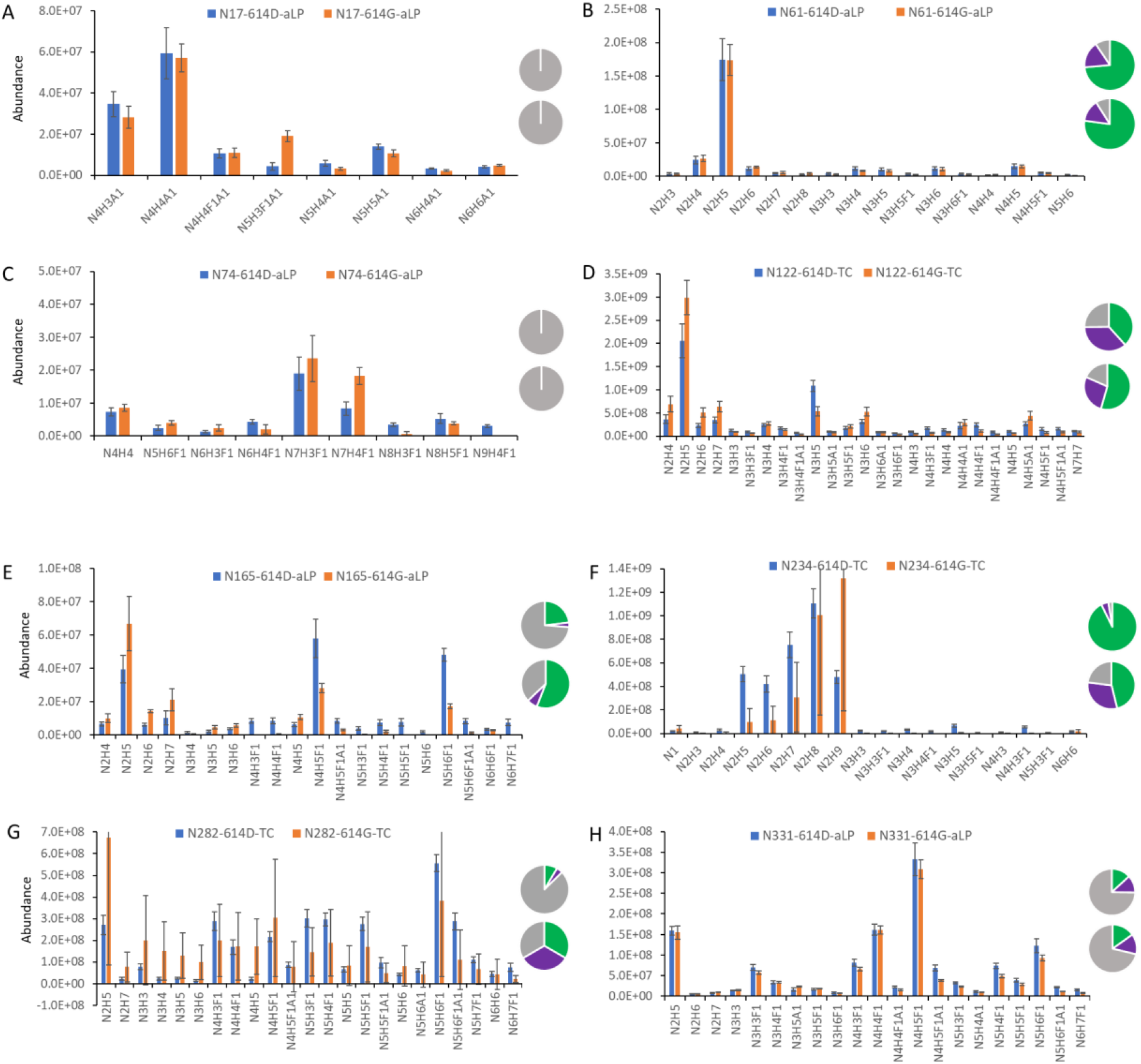
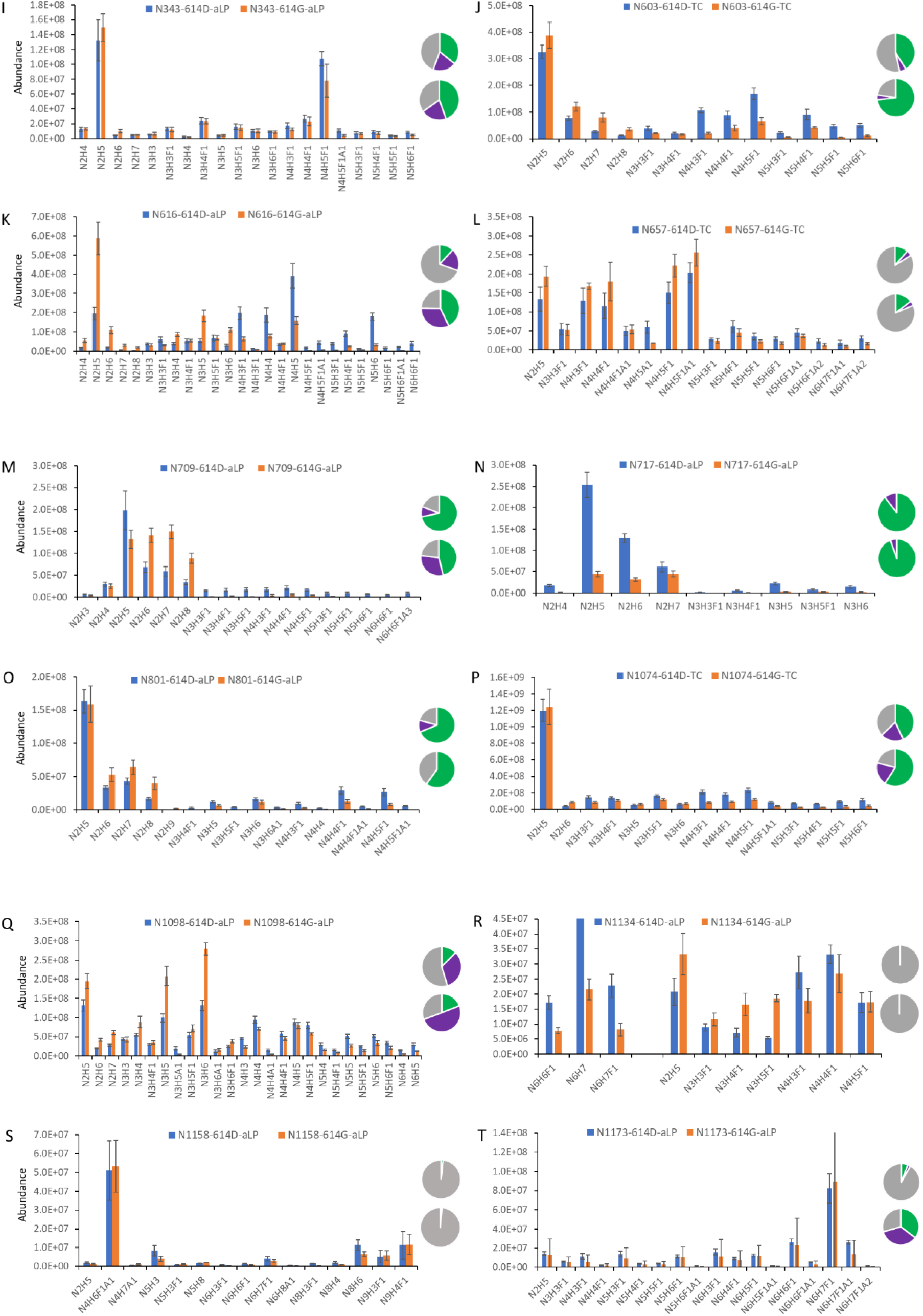
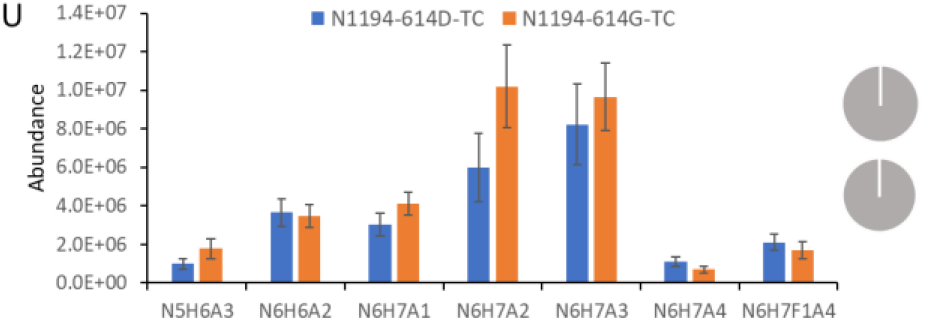
Comparison of N-linked glycosylation profiles in ectodomain spike protein between S-614D (blue) and S-614G (orange) samples. Glycosylation abundances were calculated from one representative native peptide sequence for glycan sites (A) N17, (B) N61, (C) N74, (D) N122, (E) N165, (F) N234, (G) N282, (H) N331, (I) N343, (J) N603, (K) N616, (L) N657, (M) N709, (N) N717, (O) N801, (P) N1074, (Q) N1098, (R) N1134, (S) N1158, (T) N1173, and (U) N1194. Inserted pie charts (upper: 614D; lower: 614G) depict the relative composition of high-mannose (green), hybrid (purple), and complex (gray) types of glycoforms. In the short names of individual glycans, N, H, A, and F symbolize HexNAc, Hex, NeuAc, and Fuc, respectively. X-axis represent individual glycans.

### Glycosylation sites (sequons) containing unchanged glycan profiles between two S proteins

The N-glycosylation composition on the ectodomain of the SARS-CoV-2 S-614D protein was similar with those detected on the recombinant SARS-CoV-2 S proteins by different laboratories, including ours^6, 28, 32, 37^. Analysis of the S-614G variant, however, revealed differences in the N-glycan on some glycosylation sites from those seen in S-614D. Among 21 detected and quantified sequons, 10 of these sequons including N17, N61, N74, N331, N343, N657, N1074, N1158, N1173, and N1194 had little to no significant variations in the distribution of both individual glycans and glycan types between S-614D and S-614G protein expressions while alterations were observed on 11 of sequons that include N122 N165, N234, N282, N603, N616, N709, N717, N801, N1098, N1134(Figure 2). Within each of the unchanged glycosylation sites, not only did the numbers and forms of heterogeneous glycans remain unchanged, but their relative abundances also remained unchanged.

The unchanged glycosylation sites spanned the entire surface of the S trimers - head and stalk regions, or S1 and S2 subunits (Figure 3). N1158, N1173, and N1194 are three N-glycosylation sites that reside in the stalk region or C-terminal portion of the S2 (membrane fusion subunit) subunit proximal to the viral membrane. Conserved glycosylation on these three sites observed between the two variants in our study suggests that the shielding of the stalk region by complex glycans on the S-614G variant virus remains intact.

**Figure 3.**
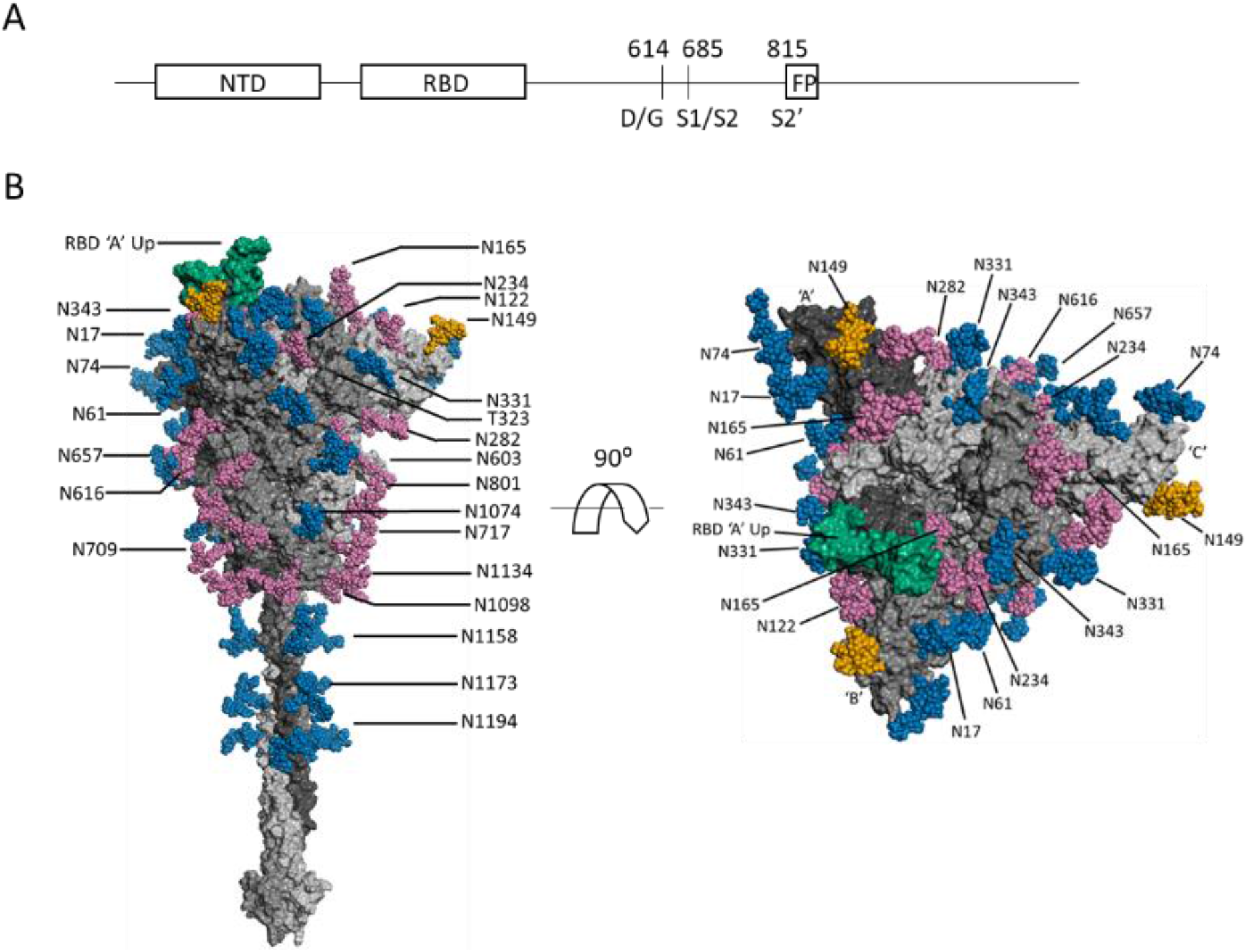
(A) Schematic SARS-CoV-2 S protein primary structure. (B) N-glycans depicted on a representative full-length, fully-glycosylated prefusion conformation of the trimetric SARS-CoV-2 spike protein (file 6vsb 1 1 1.pdb from the CHARMM-GUI Archive displayed in PyMOL)^35, 36^. Blue-colored glycans indicate no change in the glycosylation site between the S-614G mutant and the S-614D wild type. Magenta-colored glycans indicate a modification in the glycan distribution and type between the mutant and wild type. The RBD is shown in green. The N-149 glycans are gold. The glycans depicted do not necessarily match those described in this report, and the O-linked glycans in the model are hidden due to low occupancy.

Among the 7 unchanged sites in the head region, three (N17, N61, and N74) were located at the N-terminal portion of the receptor binding domain, implying that the site-specific glycosylations on this distal portion of the receptor binding domain may not be involved in enhanced binding affinity of S-614G to hACE2 receptor. N331 and N343 are the only two residues in the RBD that are modified by N-glycans. However, they are not located within the receptor binding motif of the RBD and do not directly interact with hACE2. Our analysis revealed that N-glycan microheterogeneities on each of these two sites did not change between S-614G and S-614D (Figure 2H, 2I), suggesting that the glycan complement on these sites does not affect the RBD-ACE2 binding directly or indirectly, and may only provide protection for the RBD region. The effect of the glycosylation of these two sites, if any, on any differences between the two S proteins observed in bioassays{Cerutti, 2021 #103} requires further investigation.

### Glycosylation sites with altered N-glycan profiles between S-614G and wild-type S protein

All sequons bearing glycan variations between spike protein 614D and 614G reside in the head region of the S proteins with some in the S1 subunit (N122, N165, N234, N282, N603, and N616) and others in the top half of the S2 subunit (N709, N717, N801, N1098, and N1134) at the lower portion of the head (FigFigure3). It is interesting that nearly all sequons in the lower portion of the head showed significant difference in glycan content between S-614D and 614G, in contrast to the observations in the top head area (S1 subunit) where only half of the sequons displayed a change in glycosylation forms (N1074 was the exception) (Figure 2). There might be two possible reasons for this phenomenon. One is the adaptability of the N-glycan shielding layer to the structural changes in the original protein caused by the S-D614G substitution. Another possible reason could be that the N-glycosylation in this part of the S2 subunit may play a role in viral membrane fusion, because the fusion peptide is buried in the prefusion structure, and S2 is responsible for virus-host-cell membrane fusion. Further investigation is needed to address these hypotheses.

Based on the scope of alteration, these altered glycosylation sites can be categorized into two major groups: 1) with increased high-mannose and decreased hybrid and complex glycans on the sites of N122, N234, N603, N709, and N801; 2) increased high-mannose and hybrid glycoforms and decreased complex glycans on N165, N282, N616, N1098 and N1134. Alteration on N717 showed a different trend from the other sequons. Although only glycosylated peptides were detected and the non-glycosylated N717 was not observed, the ion signals of the N717 glycopeptides were significantly reduced. This site was mainly occupied by oligomannose with little hybrid and no complex content (Table 1). Variations occurred on two major oligomannoses, Man5 and Man6, but their mass spectral abundances were reduced by approximately eight- and four-fold, respectively, when the aspartic acid at 614 was substituted for a glycine residue (Figure. 2N). Interestingly, the Cryo-EM structure of two proteins (PDB: S-614D-6vsb, 614G-6xs6) show significant differences in the secondary structures proximal to the N717 residue, which could affect the enzymatic digestion for the spike protein near this site and lead to diminished ion signals for the glycopeptides.

**Table 1.** Relative abundance (%) of the N-glycans on some of the sequons of the recombinant SARS-CoV-2 Spike D614G mutant and its ancestor.

| Group* | Sequon | Spike - 614D |  |  | Spike - 614G |  |  |  |  |  |
| --- | --- | --- | --- | --- | --- | --- | --- | --- | --- | --- |
|  |  | High-mannose | Hybrid | Complex | High-mannose | Change | Hybrid | Change | Complex | Change |
| 1 | N122 | 38.4 | 36.4 | 25.2 | 54.5 | 16.1 | 27.2 | -9.2 | 18.3 | -6.9 |
|  | N234 | 92.7 | 4.7 | 2.6 | 98.6 | 5.9 | 0.5 | -4.2 | 0.9 | -1.7 |
|  | N603 | 41.2 | 5.4 | 53.4 | 73.2 | 32.0 | 4.2 | -1.2 | 22.5 | -30.9 |
|  | N709 | 71.5 | 9.4 | 19.0 | 92.5 | 21.0 | 2.3 | -7.1 | 5.2 | -13.8 |
|  | N801 | 68.8 | 10.4 | 20.8 | 87.7 | 18.9 | 5.3 | -5.1 | 7.1 | -13.7 |
| 2 | N165 | 23.2 | 3.0 | 73.8 | 55.9 | 32.7 | 7.0 | 4.0 | 37.1 | -36.7 |
|  | N282 | 8.5 | 4.1 | 87.4 | 19.4 | 10.9 | 15.6 | 11.5 | 65.0 | -22.4 |
|  | N616 | 11.9 | 18.6 | 69.5 | 42.7 | 30.8 | 32.5 | 13.9 | 24.8 | -44.7 |
|  | N1098 | 33.6 | 33.4 | 53.6 | 19.5 | -14.1 | 50.3 | 16.9 | 30.2 | -23.4 |
|  | N1134 | 13.9 | 14.4 | 71.7 | 22.5 | 8.6 | 31.6 | 17.2 | 45.9 | -25.8 |
| 3 | N717 | 89.6 | 10.4 | 0.0 | 94.6 | 5.0 | 5.4 | -5.0 | 0.0 | 0.0 |

\* The sequons in group 1 reduce the hybrid and comple glycans in S-614G. The sequons in group 2 contain reduced complex and increased hybrid glycans in the S-614G. The N717 in the group 3 does not contain complex glycan in both S-614D and S-614G proteins.

The relative abundance of complex-type glycans at all altered N-glycan sites, (except N717, which carries no complex glycans on either S-614D or S-614G), were reduced, often significantly (i.e., 13 – 45%) (Table 1). While 6 of 11 sequons, including N165, N282, N603, N616, N1098, and N1134, were occupied by more than fifty percent of the complex glycans present on the S-614D protein, only one of them (N282) maintained 65% population of complex glycans after the S-D614G substitution. Meanwhile, the number of the sequons bearing more than 50% high-mannose glycans increased from 3 (N234, N709, and N801) to 6 (N122, N165, N234, N603, N709, and N801). This phenomenon should not be caused by lack of processing enzymes because the two proteins were expressed in identical Expi293F cells at the same time under the same conditions. The replacement of fully processed complex glycoforms with under-processed oligomannose glycans implies that less dense saccharide moieties in the S-614G protein might be sufficient to reshape the protein (or facilitate protein folding).

The closest glycosylation site to the S-D614G substitution is N616, as it resides only two amino acid residues downstream of the substitution site. Glycosylation patterns between the S-614D and S-614G proteins significantly varied at N616 (Figure 2K). This site in S-614D was occupied predominantly by several fucose-containing bi-antennary or tri-antennary complex glycans including HexNAc_4_Hex_5_Fuc_1_, HexNAc_4_Hex_3_Fuc_1_, and HexNAc_5_Hex_6_Fuc_1_ with only 12% high-mannose glycans (Figure 2K, Table 1). However, when S-614D is substituted by the smallest amino acid, glycine, the percentage of complex glycans was reduced by almost half of its original occupancy and the content of oligomannose increases by 30% and hybrid by 14%. The smallest high-mannose glycan, HexNAc_2_Hex_5_ (the high-mannose glycans were represented as Man5 – Man9 thereafter), became the most abundant glycan at N616 in S-614G protein and the MS1 peak intensities of this glycan elevated by three-fold while the intensity of the most abundant complex-type glycan, HexNAc_4_Hex_5_Fuc_1_, in S-614D decreases by more than two-fold. Although the intensity values might not reflect real changes in abundance because they were derived from two peptides, the trends of glycan distribution should not be affected significantly. Based on the Cryo-EM structures, it has been proposed that S-D614G substitution allosterically leads to more “open” conformations or a higher percentage of RBD in a “up” position that facilitates the interaction of RBD with the ACE2 receptor^13, 38^. Lower glycan complexity at the N616 site might adapt or compensate the changes in protein structure within the allosteric pathway and render the allostery effect on the mutated S protein.

Similar changes occurred on N603 (Figure 2J) and N165 (Figure 2E), where more than 30% increase in high-mannose and decrease in complex glycans were observed but the content of hybrid glycans did not change as much as the complex or high mannose structures. Considering the proximity of N603 to the S-D614G substitution site, the glycan variation on N603 might have similar effect as that on the N616.

MD simulation using a Man5 oligomannose glycan at N165 has revealed that the glycan plays an important role in modulating the conformational “Up” and “Down” transitions of the S RBD by occupying the space between the RBD and NTD regions at an “Up” position^7^. Our data showed a large decrease in the complex glycan content and an increase in oligomannose glycans with Man5 becoming the most abundant glycan in the S-614G variant. Further study is required to illustrate the effect of varied N-glycosylation on this particular site.

N234 is another glycosylation site within the NTD region that has been proposed to modulate the RBD conformational dynamics^7^. MD simulation indicates that a Man9 high-mannose N-glycan at this site fills the vacancy left by an opened RBD reaching to the apical core of the trimer and stabilizing the receptor binding domain of S protein in the “up” or “open” conformation. Our data showed that the N234 was predominantly occupied by five high-mannose glycans in both S-614D and S-614G but their distributions were different. N234 in the S S-614G was dominated by Man8 and Man9 but highest abundant glycan was Man8 in the S S-614D and four other high-mannose glycans (Man5, Man6, Man7, and Man9) had relatively similar concentrations (Figure 2F). How this change will affect the binding of RBD to human ACE2 receptor requires more investigation using biological or biophysical methods.

N709 and N801 in Group 1 are two oligomannose dominated sequons that displayed similar variation between the S-614D and S-614G proteins (Figure 2M, 2O). Whereas the combined population of complex and hybrid N-glycans declined approximately 20%, significant changes appeared within the four individual high-mannose glycans ranging from Man5 to Man8. Three of the sites had more abundant high mannose glycans in S-614G than in S-614D proteins. N122 is the last member in Group 1 with elevated high-mannose and reduced hybrid and complex glycans. The main changes occur on one of the high-mannose glycan (Man5) where its abundance in S-614G increased by 50% in the S mutant and was much higher than that of the other glycans (Figure 2D). Because these two residues are located downstream from the S1/S2 cleavage site, reduced complexity of their micro-heterogeneous glycoforms might be related to enhanced protease susceptibility at the furin cleavage site determined by bioassays^38^.

In addition to N165 and N616, Group 2 includes three other sequons, N282, N1098 and N1134, with the sites originally occupied predominantly by highly processed complex-type in the S-614D proteins, suggesting that further processing beyond the high-mannose forms was favored on these sites. Unlike the members in Group 1, reduced populations of complex glycans were compensated by increased hybrid populations (Table 1). In contrast to some sites where a few glycoforms dominated the occupancy, the abundances of various glycoforms were distributed more evenly in both S-614D and S-614G S proteins (Figure 2G, 2Q, 2R). Since N1098 and N1134 are located in the C-terminal S2 domain and far from the RBD and NTD domains, the glycosylation of these sites might not affect interactions between spike’s RBD and human ACE2 receptors.

## Conclusion

We analyzed the N-glycosylation profiles of the SARS-CoV-2 spike D614G variant and its ancestor protein using advanced EThcD mass spectrometry. To provide accurate comparison, two recombinant proteins were prepared using identical cells and under the same experimental conditions. The results of this study revealed that the single D614G substitution had impacts on the glycosylation of the spike protein and half of the N-glycosylation sequons in the S showed a difference in the distribution of various glycan forms between the S-614D and S-614G variants. The relative abundances of the complex-type glycans were reduced by up to 45% on those sequons with altered N-glycosylation in the S-614G glycoprotein and the contents of high-mannose glycans, in contrast, were elevated by up to 33%. This data shows reduced overall complexity of the N-glycosylation on the S-614G protein, which likely play a role in reshaping the homotrimer. Our findings also included the variation of the N-glycan forms on the two major parts of the ectodomain spike, head, and stalk. It was observed that the all the glycosylation sites displaying different modification profiles between the two proteins were located in the spike head while glycosylation of three sites in the stalk showed the same N-glycosylation pattern for both the S-614D and S-614G and glycoproteins. These results provide important information toward understanding the effects of protein glycosylation on the structure and function of the SARS-CoV-2 spike protein.

## Methods

### Materials

All chemicals were obtained from Sigma–Aldrich (St. Louis, MO) except where otherwise indicated. Endoproteases including trypsin and chymotrypsin were purchased from Promega (Madison, WI), and α-lytic protease was obtained from New England BioLabs (Ipswich, MA). The recombinant poly-histidine tagged ectodomains of the SARS-CoV-2 S proteins (614D and 614G) were expressed in human Expi293F cells using the Expi293 Expression System (ThermoFisher Scientific) and purified using HisTrap FF column (GE Life Sciences)^18^. The sequences of the two proteins include mutated furin cleavage site and K986P, V987P substitutions. Biotinylated ACE2 was purchased from Sino Biological Inc (Wayne, PA). Mouse SARS-CoV2 monoclonal antibodies (3G7 and 3A2) against the RBD of the spike were prepared by an accelerated immunization and hydridoma screening process^33^

### Binding assay and 1D Gel electrophoresis

For binding assays, the biotinylated ACE2 was immobilized to Streptavidin magnetic beads and the monoclonal antibodies (mAb) were attached to Protein G magnetic beads. A mixture of 1 μg of S protein and equal amount of ACE2 on beads or mAb on beads in 0.6 mL phosphate-buffered saline with 0.05% Tween 20 was incubated with gentle rotation for 60 min. Sodium dodecyl sulfate–polyacrylamide gel electrophoresis (SDS-PAGE) was performed on a NuPAGE Novex Bis-Tris gel following manufacturer’s instructions (Invitrogen). Protein/ACE2 or protein/mAb attached on magnetic beads or protein control in solution were mixed with 4X Sample Buffer and deionized water (1:3 v/v) and heated at 80 °C for 10 min. The supernatants were loaded on a 4–12% gradient gel, and the gel was run in MOPS buffer at 200 V for 45 min. The gels were stained with the SYPRO Ruby Protein Gel Stain (Thermo Scientific, Waltham, MA).

### Digestion of spike proteins

The enzymatic digestion of recombinant proteins were conducted as previously reported with modifications^32^. The aliquots of full-length ectodomain spike protein ( S-614D or 614G) (1 μg) were denatured and reduced at 90°C for 20 min in a solution containing 50 mM ammonium bicarbonate (pH 7.8), 0.015% RapiGest SF surfactant (Waters Corporation) and 1.5 mM DTT. Samples were alkylated using iodoacetamide for 30 minutes in the dark at room temperature with gentle mixing. Six identical aliquots of each protein were prepared for two digestion methods, and each digestion was performed with three replicates. The digestions with α-lytic protease were conducted at 37°C overnight at an enzyme to protein ratio of 1:10 (w/w). For the samples digested sequentially by two enzymes, the first digestion was performed at 52°C for 60 min with trypsin (1:3 w/w) and the second digestion was conducted at 37°C overnight using chymotrypsin at an enzyme-to-substrate ratio of 1:15 (w/w). The proteolytic reactions were quenched and the RapiGest was precipitated by adding 5% trifluoroacetic acid to decrease the pH to below 3. The mixture was then incubated at 37°C for 30 min. The solutions were centrifuged at 4000 RPM for 10 min, and the supernatants (20 μL) were transferred into new sample vials.

### Mass spectrometry analysis

Nanoflow liquid chromatography coupled to electrospray ionization tandem-mass spectrometry (LC-MS/MS) analysis was performed on an Orbitrap Eclipse Tribrid mass spectrometer and an UltiMate^™^ 3000 RSLCnano chromatography system (Thermo Scientific) as described previously^32^. The protein digest was separated on an integrated separation column/nanospray device (Thermo Scientific EASY-Spray PepMap RSLC C_18_, 75 μm i.d. x 15 cm length, 3μm 100Å particles), coupled to an EASY-Spray ion source. The mobile phase was 0.1% formic acid in water (mobile phase A) and 0.1% formic acid in 80% acetonitrile/20% water (mobile phase B) using the following gradient: 4% B (0 – 8 min); 4 – 10% B (8 – 10 min); 10 – 35% B (10 – 43 min); 35 – 60 % B (43 – 45 min); 60 – 95% B (45 – 46 min); 95% B (46 – 53 min); 95 – 4% B (53 – 53 min); 4% B (53 – 63 min). The flow rate was 300 nL/min and 8 μL of each sample was injected in duplicates. The spray voltage was set to 1.8 kV, and the temperature of the integrated column/nanospray device was set at 55°C. The temperature of the ion transfer tube was set to 275°C.

Mass spectrometric data acquisition was performed using a signature ion triggered electron-transfer/higher-energy collision dissociation (EThcD) method^34^. MS precursor scans were acquired by the orbitrap at a resolution of 120,000 (measured at *m/z* 200), from *m/z* 375 – 2000 with the automatic gain control (AGC) target setting as “standard” and the maximum injection time as “auto”. An initial data-dependent MS/MS scan was acquired using HCD at a resolution of 30,000, a mass range of *m/z* 120 – 2000, and a normalized collision energy (NCE) of 28%. Signature ions representing glycan oxonium fragments were used to trigger the ETD fragmentation. If one of three common glycan signature ions, *m/z* 204.0867 (HexNAc), 138.0545 (HexNAc fragment) or 366.1396 (HexNAcHex), was detected in the HCD spectrum within 15 ppm mass accuracy, an additional precursor isolation and EThcD acquisition was performed at a resolution of 50,000 (measured at *m/z* 200), scan range of *m/z* 150 – 2000 with normalized AGC Target = 500% and maximum injection time = 150 ms. The supplemental activation NCE was set to 35%.

### Data analysis

MS/MS data were processed using PMi-Byonic (Version 3.7) node within Proteome Discover (Thermo Scientific) Data were searched using the Protein Metrics 182 human N-glycan library (included in the Byonic program) as potential glycan modifications. The search parameters for enzyme digestion were set to fully specific, 3 allowed missed cleavage sites, and 6 ppm and 20 ppm mass tolerance for precursors and fragment ions, respectively. Carbamidomethylation of cysteine was set as a fixed modification with variable modifications set to include deamidation at Asn and Gln and oxidation of Met. Tandem mass spectra of identified glycopeptides with a Byonic^™^ score higher than 150 were considered valid identifications. Identified glycopeptide and unoccupied peptide abundances were determined using precursor ion peak intensity with normalization on total peptide amount per file. Relative abundance of each type of glycans at each site was calculated as the normalized peak intensity ratio of the peptides bearing a particular glycan type over the total glycopeptides. The glycan abundance was represented as the mean of six replicates along with standard deviation of the mean. The glycan abundances of several glycosylation sites including N122, N603, N616, N717, N1074, and N1158 was summed from the average values of two peptides bearing an identical sequon.

### Representative 3D structure model

A PDB file containing a complete model of the full-length fully-glycosylated S trimer was obtained from the CHARMM-GUI Archive^35, 36^. The file 6vsb_1_1_1.pdb was chosen as being representative of S trimer with one chain in the “up” position. The model, based on the partial PDB: 6VSB cryo-EM structure, incorporates wild-type residues 1-1273 with 22 N-linked glycans and 1 O-linked glycan per monomer [1]. The model was displayed using PyMOL, the only change being that the O-linked glycans were hidden due to low occupancy [3]. The displayed glycans do not necessarily match those described in this paper.

## Data availability

The datasets generated and analyzed in the scope of this study are available from the corresponding authors upon request.

## Author contributions

D.W., J.R.B and J.L.P. designed the study. D.W. did the measurements, analyzed the data, prepared figures and tables, and wrote the manuscript. T.K., M.S., J.B., J.L.B. and S.O. processed the data. A.R.W. prepared the figure. B.Z., X.F., and D.E.W. prepared recombinant spike proteins. J.G., M.G.F., A.P. C. provided antibodies. All authors conceived the study and reviewed the manuscript.

